# Activation of basal forebrain cholinergic neurons promotes the attribution of incentive salience to reward cues in male rats

**DOI:** 10.64898/2026.10.20.760494

**Authors:** Elizabeth A. Rakowski, Christopher P. King, Ashmita Mukherjee, Esther Ruhl, Luke Hannan, Gregory C. Loney, Paul J. Meyer

## Abstract

**Background:** Reward-associated environmental stimuli (“cues”) can gain incentive salience in some individuals, thereby becoming attractive, reinforcing, and instigators of motivational states. In rodents, the tendency to attribute incentive salience can be measured using a Pavlovian Conditioned Approach (PavCA) procedure, in which approach to a food cue (‘sign-tracking’), relative to approach to the food delivery location (‘goal-tracking’) is the operational definition of the incentive salience attributed to the cue. Growing evidence suggests that acetylcholine promotes this process, thus the current study tested the effect of activating cholinergic neurons in the basal forebrain on sign- and goal-tracking during PavCA.

**Methods:** We injected a Cre-dependent virus with excitatory Designer Receptors Exclusively Activated by Designer Drugs (DREADDs) into the horizontal limb of the diagonal band of Broca of transgenic rats expressing cre-recombinase in acetylcholine containing neurons. We then injected clozapine-n-oxide (CNO) before each of eight sessions of PavCA. On the ninth session, no CNO was given to test if any effects were immediately reversible.

**Results:** We found that activation of basal forebrain cholinergic neurons increased sign-tracking over time without an overall effect on goal-tracking, relative to a control group injected with a control virus. Changes in responding from the final CNO session to the saline test did not differ between virus groups.

**Conclusions:** This effect demonstrates that acetylcholine promotes the attribution of incentive salience to a reward-related cue. Future studies should investigate the effect of inhibiting this area on sign-tracking as well as how activation of this area influences reactivity to cues associated with other rewards such as drugs of abuse.

## INTRODUCTION

Motivated behavior is influenced by rewards (unconditioned stimuli, **US**) as well as the cues associated with these rewards (conditioned stimuli, **CS**). Through Pavlovian learning, cues can acquire incentive salience, as measured by the ability to elicit approach, reinforce new actions, and trigger motivational states (Milton & Everitt, 2010; Robinson et al., 2014). Importantly, while cues predict rewards for all individuals, some differ in the degree to which incentive salience is attributed to those cues. In preclinical studies using a Pavlovian conditioned approach (**PavCA**) procedure, animals that tend to approach and interact with the reward-related cue are known as “sign-trackers” while other animals that approach the reward delivery site when the cue is present are known as “goal-trackers” (Meyer, Lovic, et al., 2012; Tomie et al., 2000; Flagel et al., 2007; Boakes, 1977; Hearst and Jenkins, 1974). In conditioned reinforcement, sign-trackers will work to obtain the cue even when the reward is absent, indicating that the cue has acquired motivational properties beyond simple prediction (Meyer, Lovic, et al., 2012; Robinson & Flagel, 2009). Sign-tracking can also be relatively resistant to outcome devaluation, suggesting that cue-directed behavior may persist despite changes in reward value (Morrison et al., 2015). This individual propensity to attribute incentive salience to reward cues is influenced by genetic factors (King et al., 2025) and is particularly relevant to substance use disorder (**SUD**), where drug-paired cues can trigger craving and relapse. For example, sign-trackers exhibit a variety of addiction-relevant behaviors, including: approach to Pavlovian drug cues (Yager et al., 2015; Yager & Robinson, 2013, 2015), sensitivity to psychostimulant cues during intravenous self-administration (IVSA) (Beckmann et al., 2011; Everett et al., 2020; Kawa et al., 2016; Saunders & Robinson, 2010; Tunstall & Kearns, 2015), nicotine IVSA and cue-induced reinstatement (Versaggi et al., 2016), cocaine-induced ultrasonic vocalizations (Meyer, Ma, et al., 2012; Meyer & Tripi, 2018; Tripi et al., 2017), alcohol-directed behaviors (Tomie et al., 2004; Tomie et al., 2002), resistance to punishment during remifentanil seeking (Zumbusch et al., 2023), and impulsivity (King et al., 2016; Lovic et al., 2011).

These behavioral traits are subserved by neurobiological processes that also differ between sign- and goal-trackers. Sign-trackers exhibit more excitation in brain areas related to motivation such as orbitofrontal cortex and nucleus accumbens (Flagel, Clark, et al., 2011; Moorman & Aston-Jones, 2014). During Pavlovian conditioning for a food reward, sign-tracking was correlated with higher dopamine-1 receptor mRNA expression in the nucleus accumbens and sensitivity to dopamine receptor agonists (Flagel et al., 2007; Amissah et al., 2026; Roughley and Killcross 2019; Fraser et al., 2016). Dopamine antagonism in the nucleus accumbens core prevents the acquisition of sign-tracking but not goal-tracking (Flagel, Clark, et al., 2011; Danna and Elmer, 2010) and sign-trackers exhibited more cue-elicited neuronal activation in reward-related areas like the nucleus accumbens, ventral pallidum, and caudate putamen, implying that the incentive value of a cue is a crucial component in motivated behaviors (Flagel, Cameron, et al., 2011; Saunders et al., 2018; Ahrens et al., 2016; Gillis and Morrison 2019; Duffer et al., 2023).

Acetylcholine (ACh) is critical for attentional processes that establish and maintain the relationship between rewards and their cues (Sarter and Lustig, 2019). ACh is generally involved in managing both “top-down” and “bottom-up” sensory processing, where the former mechanism discriminates specific stimuli from “noise” and the latter sends information about salient stimuli from the environment to cortical areas responsible for more complex processing (Bentley et al., 2004; Sarter et al., 2001). Cholinergic neurons in the basal forebrain act as modulators of attention to reward-related cues, as evidenced by an increase in cholinergic activity originating from this area during correct cue detection (Gritton et al., 2016; Parikh et al., 2007; Sarter et al., 2005) and a decrease in correct performance in attention tasks when this area is inhibited or lesioned (Chiba et al., 1999; McGaughy et al., 1996; Muir et al., 1994).

Differences in cholinergic systems likely play a role in the attribution of incentive salience to reward cues. Stimulation of acetylcholine receptors promotes sign-tracking, while blockade of these receptors inhibits it (Versaggi et al., 2016; Palmatier et al., 2013; Overby et al., 2018; Gheidi et al., 2023; King and Meyer 2022). Paradoxically, sign-trackers have deficits in cholinergic signaling that result in attentional impairments and may relate to their addiction-related phenotypes (Paolone et al., 2013; Pitchers et al., 2017; Phillips and Sarter, 2020). Further, chemogenetic inhibition of the basal forebrain, especially the horizontal limb of the diagonal band (**HDB**), impaired attention in goal-trackers, but had no effect in sign-trackers (Kucinski et al., 2022). These findings implicate basal forebrain projection systems in the attentional control associated with goal-tracking, and that this system may be compromised in sign-trackers.

Thus, there is a paradox where sign-trackers have deficits in some aspects of the cholinergic system, but stimulation of cholinergic receptors pharmacologically enhances sign-tracking. This may be due to the contrasting roles of acetylcholine in attention and incentive salience attribution. However, whether stimulation of basal forebrain cholinergic neurons enhances sign-tracking during Pavlovian conditioning remains unclear. Therefore, in this study we used a chemogenetic approach to activate these neurons and determine the effect of this activation on sign- and goal-tracking behavior. Given the previous findings using acetylcholine receptor agonists, we hypothesized that activation of cholinergic neurons in the basal forebrain would increase sign-tracking during a Pavlovian-conditioned approach paradigm, with no effect on goal-tracking behavior.

## METHODS

### Animals

Fig. 1 shows a brief timeline of the experiment. Male ChAT-Cre rats (final n=52) aged 10 to 12 weeks were bred at the University at Buffalo Psychology Department from breeders obtained from the Rat Resource & Research Center (LE-Tg(Chat-Cre)5.1; Deis https://www.rrrc.us/; Witten et al. 2011). We chose male rats to avoid ceiling effects when using females, who sign-track at higher rates than males (King et al., 2021). These rats, created on a Long-Evans background, produce cre-recombinase only in neurons expressing choline acetyltransferase (**ChAT**). Only heterozygous, cre-positive rats were used and were identified using tail or ear samples by Transnetyx (Cordova, TN). Rats were pair-housed in standard cages with a reverse 12-hour light cycle with *ad libitum* access to food and water. They were handled for five to seven days before experimentation began. All experimental procedures were approved by the University at Buffalo Institutional Animal Care and Use Committee and followed the Guide for the Care and Use of Laboratory Animals (National Research Council, 2011).

**FIGURE 1:**
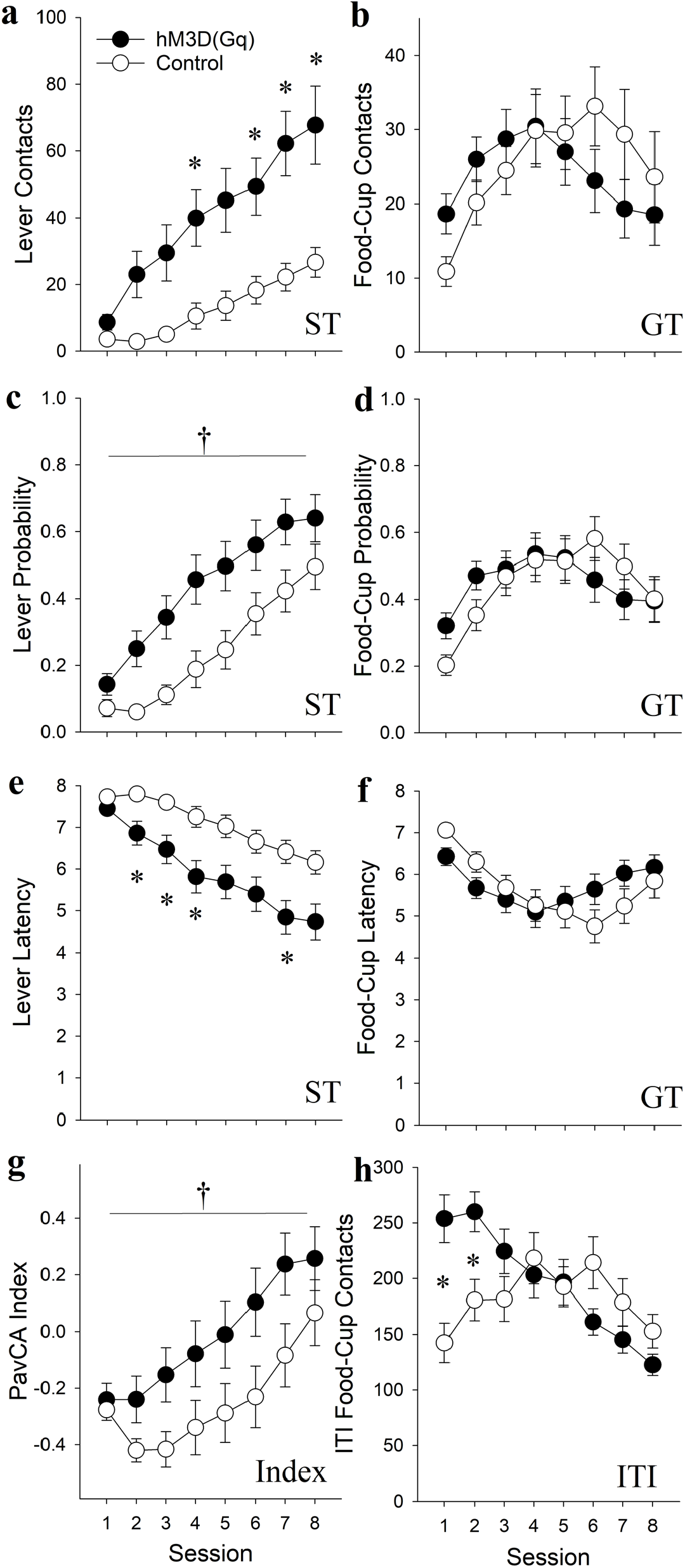
Stimulation of cholinergic neurons in the basal forebrain enhanced sign-tracking without overall effects on goal-tracking behavior. Timeline (*above*) shows the sequence of experiment events. Panels (a), (c), and (e) show measures of lever-directed approach (sign-tracking, ST), while panels (b), (d), and (f) show food-cup oriented approach (goal-tracking, GT). Panel (g) shows the Pavlovian conditioned approach (PavCA) index scores, while panel (h) shows inter trial interval (ITI) food-cup entries. Asterisks indicate between-groups differences on specific sessions using a Bonferroni-adjusted alpha of .000625. Daggers indicate a main effect of *Virus* (p<0.05).

### Stereotaxic Surgery

To activate cholinergic neurons, we used a chemogenetic approach using viral-mediated transfer of hM3D(Gq), an excitatory designer receptor exclusively activated by designer drugs (**DREADDs**; Armbruster et al. 2007; Ferguson and Neumaier, 2012; Smith et al., 2016). The experimental group was injected with a cre-dependent construct into the basal forebrain of ChAT-cre rats [pAAV-hSyn-DIO-hM3D(Gq)-mCherry; Addgene #44361-AAV5], thereby inducing the expression of DREADDs specifically in cholinergic neurons. A control group received a control construct [AAV-hSyn-DIO-mCherry; Addgene viral prep #50459-AAV5], which contained only the mCherry fluorescent marker used to confirm placement using immunohistochemistry. The final sample included 52 rats, with 28 in the hM3D(Gq) group and 24 in the mCherry control group.

These viral constructs were delivered using standard stereotaxic techniques under 3% isoflurane anesthesia (*e*.*g*., Wakabayashi et al., 2021; Pasquariello et al., 2018), using bupivacaine (2.5 mg/mL; 0.1 mL) and septocaine (40 mg/mL articaine HCl with epinephrine 1:100,000) as local anesthetics. The virus was injected bilaterally with a Hamilton (Reno, NV) syringe at an infusion rate of 0.2 µL/min with a total volume of 1 µL on each side. We targeted the horizontal limb of the diagonal band of Broca (HDB) based on previous studies examining cholinergic nuclei in this region (*e*.*g*. Kucinski et al., 2022; Avila and Lin, 2014). Our coordinates (AP: -0.6, ML: ±2.25, DV; -8.6 from skull) were based on our pilot studies and the Paxinos and Watson rat brain atlas (Paxinos and Watson, 2014). The needle was left in place for five minutes on each side to allow for dispersal. Carprofen (5 mg/kg) was injected subcutaneously immediately after surgery and for two days after for post-operative analgesic treatment. The animals remained in their home cages for five weeks to allow for viral expression.

### Pavlovian Conditioned Approach (PavCA)

Rats were given 45 mg banana-flavored food pellets (one tablespoon of ∼160 pellets; #F0059; Bio-Serv, Flemington, NJ) in their homecages for each of two days preceding the experiment, to minimize neophobia during testing. Standard operant chambers (20.5×24.1×29.2 cm; Med Associates Inc.; St. Albans, VT) were equipped with a food-cup attached to an automatic pellet dispenser and two LED-backlit levers adjacent to either side of the food-cup, which was located on the central panel of one wall of the chamber. A red houselight was located near the top of the wall opposite the food-cup. Rats underwent one food-cup training session the day before Pavlovian conditioning began. After a 5-min acclimation period, the houselight was turned on and 25 banana-flavored food pellets were delivered 30-second variable-interval schedule (0-60 seconds), thus the session lasted 17.5 minutes on average.

After food-cup training, rats were tested during 8 daily sessions of PavCA. All rats received an intraperitoneal injection of 1 mg/kg clozapine-N-oxide (**CNO**), the DREADD agonist, 30 minutes prior to each conditioning session. Each session consisted of a one-minute acclimation period followed by 25 trials in which an illuminated lever was inserted into the chamber for eight seconds and a food pellet was delivered upon lever retraction. These trials occurred on a 90-second (30-150 seconds) variable-interval schedule as the food-cup training with an average session duration lasting 38.5 minutes. The position of the lever-cue was assigned to the left or right side in a counterbalanced fashion. Finally, during the last session, rats received a saline injection instead of CNO to test whether any effects would be maintained upon CNO removal.

Primary measures included lever contacts, as defined by incidental lever deflections, and food-cup contacts during the CS, as defined by interruption of the photoelectric beam located within the food cup. In addition to *contacts*, we calculated the *probability* of contacting the lever or the food cup during the CS period on each trial (defined as the number of trials with a lever press or food cup entry, divided by the total number of trials). We also calculated the *latency* to the first lever deflection or food cup entry. Trials without a response during the eight-second CS were assigned a latency of eight seconds, and trial latencies were averaged to obtain the session value. Finally, we also measured the food cup contacts during the intertrial interval (**ITI**), including the interval before the first CS and the 20-second interval after the final CS. In addition to these measures, the tendency to sign- or goal-track can also be captured using the PavCA Index, as described by Meyer et al. (2012). Briefly, the PavCA Index, ranging from -1.0 to 1.0, is calculated from the average of three factors per session; *response bias* [(lever presses minus CS food cup entries) / (lever presses plus food cup entries)], *approach probability difference* [(# of trials with at least one lever press minus # of trials with at least one food cup entry during the CS) / 25 trials], and *latency difference* [(latency to enter the food cup during the CS minus latency to lever press) / 8].

### Immunohistochemistry

Rats were euthanized with 150 mg/kg sodium pentobarbital and transcardially perfused with 4% paraformaldehyde (PFA). Brains were removed and soaked in PFA for 24 hours at 4ºC and then cryoprotected with 0.1 M phosphate-buffered saline (PBS) solution with 30% sucrose and 0.1% sodium azide. Brains were coronally sectioned into 40 µm slices using a microtome (Hacker Instruments, Winnsboro, South Carolina) and stored in 0.1% sodium azide/0.1 M PBS solution at 4ºC. For immunohistochemistry, slices were rinsed in 0.1 M PBS, followed by a 0.3% Triton-X and 3% normal goat serum blocking solution. Then, slices were incubated in primary anti-RFP antibody (Rockland 600-401-379; Limerick, PA) overnight at 4ºC. Slices were then rinsed with 0.1 M PBS washes and blocking solution before incubating in secondary CF-555 (Biotium 20232; Fremont, CA) antibody for 2 hours at room temperature. Slices were mounted on slides and imaged with an epifluorescence microscope equipped with a digital camera (Echo Revolve; San Diego, CA) to verify placement of the virus. Rats with incorrect placement or no evidence of viral expression were removed from analysis (n=8).

### Statistical Analyses

Differences in approach measures during PavCA were analyzed using repeated-measures analyses of variance (**ANOVA**) with *Virus* [hM3D(Gq) vs. Control] as the between-groups variable and *Session* as the within-subjects variable. When main effects or interactions were found, Bonferroni-corrected planned comparisons were used to compare specific group differences. Independent-samples t-tests were also conducted to identify any acute changes that occurred during the CNO removal session. All statistical analyses were done using Statistica 14.1 software (Dell Inc., Tulsa, OK) and graphical representations were made using SigmaPlot 12.5 software (Grafiti LLC, St. Palo Alto, CA).

## RESULTS

Two rats were removed from testing because they did not eat the food pellets during testing. For sign-tracking measures (**Fig. 1a, 1c, 1e**), there was a significant *Virus* x *Session* interaction for lever contacts (*F*(7,350) = 3.36, *p* < .01), lever latency (*F*_(7,350)_ = 2.18, *p* < .05), and a main effect of *Virus* on lever probability (F(1,50) = 7.91, p < .01), indicating enhanced sign-tracking in hM3D(Gq) rats. There were also main effects of Virus on lever contacts (F(1,50) = 12.63, p < .001) and lever latency (F(1,50) = 10.88, p < .01). Bonferroni-corrected planned comparisons revealed that hM3D(Gq) rats made more lever contacts with shorter latency to approach the lever compared to Controls on sessions 4, 6, 7, and 8 and approached the lever with shorter latencies on sessions 2, 3, 4, and 7 (all p < .00625).

For goal-tracking measures (**Fig. 1b, 1d, 1f**), there were no main effects of *Virus*, although there were significant *Virus x Session* interactions on contacts, probability, and latency (F_(7,350)_ = 2.52-3.34, *p*’s < .05), but there were no between groups differences on any individual session. PavCA Index scores were higher in hM3D(Gq) rats across sessions compared to Controls (F(1,50) = 4.05, p < .05; **Fig. 1g**), consistent with a greater tendency to sign-track. Finally, there was a *Virus* x *Session* interaction on inter-trial food-cup contacts (*F*_(7,350)_ = 5.90, *p* < .001), where hM3D(Gq) activation caused more interaction with the food-cup on sessions 1 and 2 (*p*< .05; **Fig. 1h**). There were no statistically significant changes for the above measures during the CNO removal test (**Fig. 2**).

**FIGURE 2:**
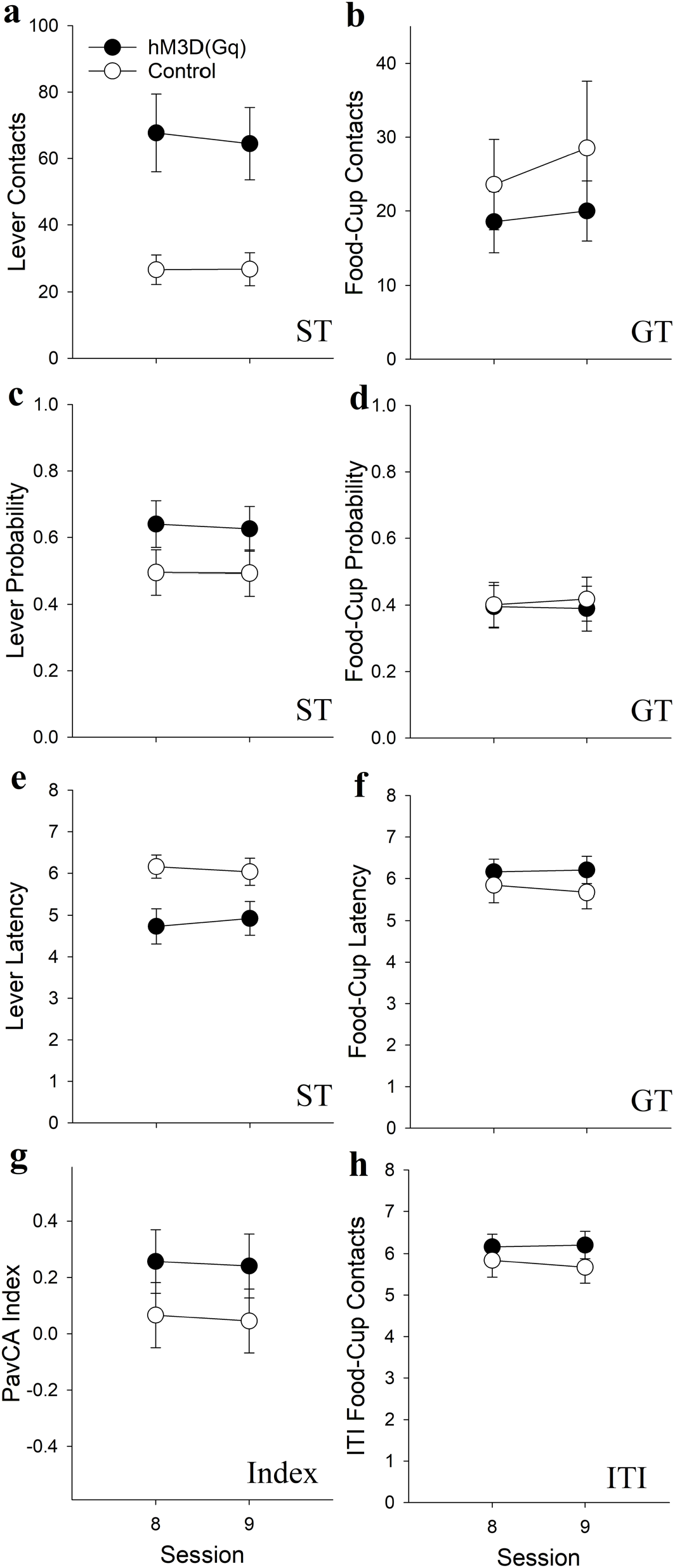
Removal of cholinergic stimulation did not affect approach behaviors. Panels (a), (c), and (e) show measures of lever-directed approach (sign-tracking, ST), while panels (b), (d), and (f) show food-cup oriented approach (goal-tracking, GT). Panel (g) shows the Pavlovian conditioned approach (PavCA) index scores, while panel (h) shows inter trial interval (ITI) food-cup entries. Data are means ± SEM; hM3D(Gq), n = 28; Control, n = 24. For each measure, the change from Day 8 to the saline session was compared between virus groups. None of these comparisons was significant (all ps > .21).

Immunohistochemistry confirmed mCherry expression in the basal forebrain (**Fig. 3**). Although injections targeted the horizontal limb of the diagonal band (HDB), expression also extended into nearby regions, including the substantia innominata, lateral and magnocellular preoptic nuclei, and ventral pallidum.

**FIGURE 3.**
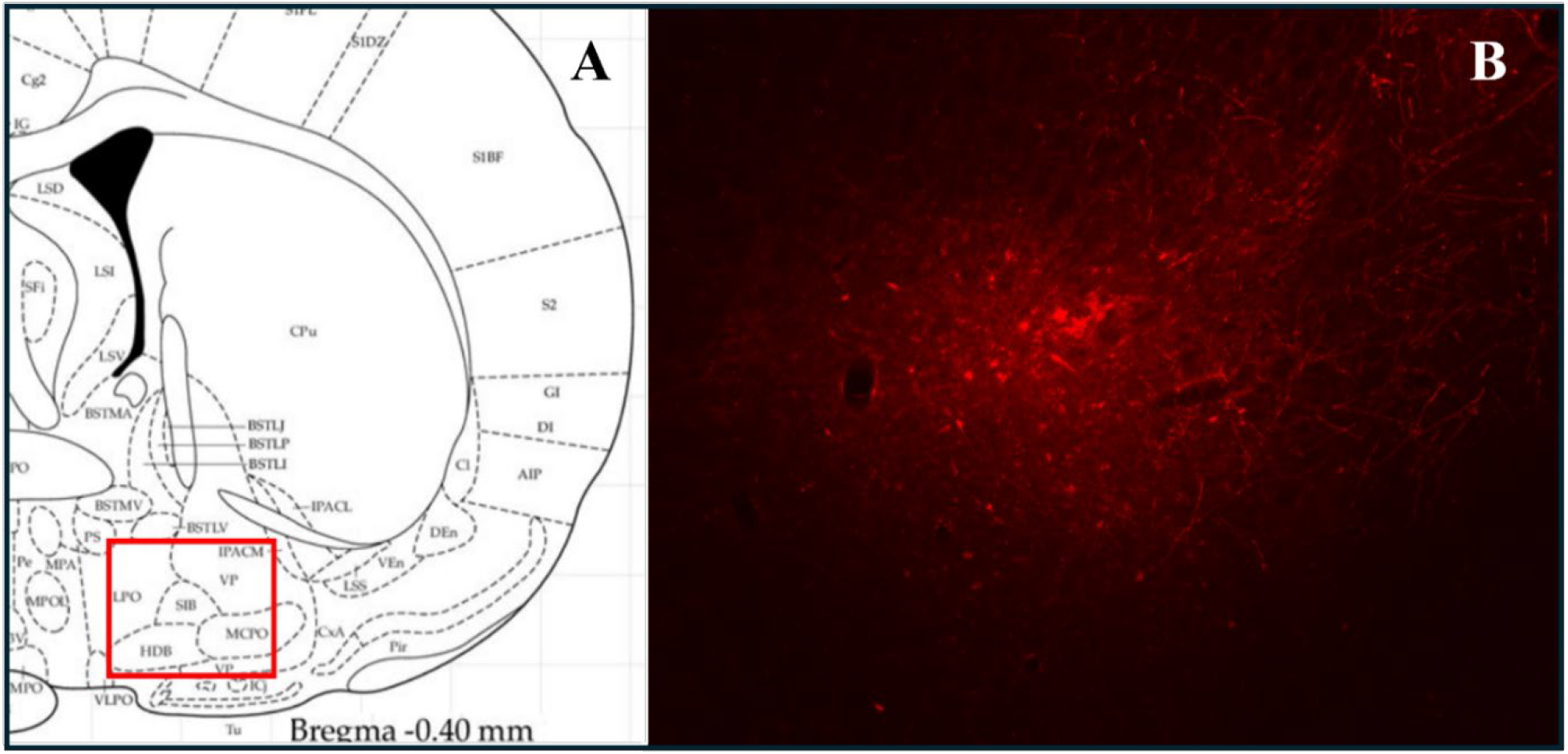
Panel A: Stereotaxic injections were targeted to the horizonal limb of the diagonal band (HDB), but expression was seen in nearby cholinergic areas (Panel B), including the substantia innominata (SIB), the preoptic nuclei (LPO and MCPO), and the ventral pallidum (VP).

## DISCUSSION

To test the effect of stimulation of cholinergic neurons in the basal forebrain on the tendency to attribute incentive salience to food cues, we tested rats in a Pavlovian conditioned approach paradigm after stimulating excitatory DREADD receptors delivered to cholinergic neurons in ChAT-cre rats. We found that activation of these neurons led to faster and more vigorous approach to the lever cue without systematic differences in approach to the food-cup. This enhancement of sign-tracking behaviors without a significant decrease in goal-tracking supports our hypothesis that basal forebrain cholinergic neurons play a part in salience attribution to reward-related cues. This is consistent with pharmacological studies showing that nicotinic receptor stimulation enhances sign-tracking in several paradigms (Palmatier et al., 2013; Versaggi et al., 2016; Ruffolo et al., 2022; Stringfield et al., 2019; Overby et al., 2018, Loney et al., 2019; but see Angelyn et al., 2021), and may play a key role in the effects of nicotine on the self-administration of opioids and other drugs (Loney et al., 2021)

However, these results appear inconsistent with findings from neurochemical studies conducted in sign- and goal-tracking rats. For example, studies on the role of cortical cholinergic signaling in attentional control have found that sign-trackers have diminished top-down attentional control in a sustained attention task (**SAT**), in which rats must detect the presence or absence of a cue in order to receive a reinforcer (Paolone et al., 2013). Sign-trackers also had blunted cholinergic tone during the task, thus linking poor attentional control with low cortical cholinergic tone. However, unlike previous studies of sign-tracking measures, the SAT measures of attention were not improved by nicotine in this study, although they were enhanced by the α4β2-contaning nicotinic receptor partial agonist ABT-089. These findings show the importance of distinguishing pharmacological effects of sign-tracking behavior from attentional traits associated with the sign-tracker phenotype. That is, a manipulation that increases sign-tracking does not necessarily reproduce the broader pattern of attentional deficits observed in sign-trackers. For example, environmental enrichment increased sign-tracking while reducing premature responding (Ishiwari et al., 2024), and the properties of the cue can substantially affect their acquisition of incentive salience (Meyer et al., 2012).

Yet, in another study, an inhibitor of the choline transporter (**CHT**), which reduced potassium-stimulated acetylcholine measured via microdialysis, also increased sign-tracking (Koshy Cherian et al., 2017). Thus, both DREADD stimulation in the current study and CHT inhibition enhanced sign-tracking, despite their different effects on cholinergic signaling.

However, these manipulations differ in mechanism and anatomical specificity and may not produce equivalent, opposing changes in acetylcholine release. Differences in phasic versus tonic levels of acetylcholine may also play a role. Generally speaking, phasic acetylcholine release is critical for cue detection (Howe et al., 2017), while tonic levels influence sustained attention and cognitive control (Parikh and Sarter, 2008). DREADD activation may increase tonic activity more than phasic bursts. This distinction could help reconcile the paradox of enhanced sign-tracking in rats with reduced cholinergic-mediated attention.

Heterogeneity within the basal forebrain is also a key consideration regarding our results. We targeted the HDB, but our virus likely transduced cholinergic neurons in other areas such as the substantia innominata (SI), the magnocellular preoptic nucleus (**MCPO**), and the ventral pallidum (**VP**). The projection targets of these areas are diverse and includes cortical areas, the hippocampus, the amygdala, and the olfactory bulb, among others (Gritti et al., 2003; Záborszky et al., 2018). The VP is of particular interest with regard to sign-tracking because these neurons are activated selectively during the presentation of reward cues (Tindell et al., 2004), and inhibition of VP neurons using inhibitory DREADDs delivered to this region reduced the acquisition of sign-tracking, and this was associated with cue-induced neural activity within this region (Chang et al., 2015; Ahrens et al., 2016). This suggests a critical role for the VP in the acquisition of sign-tracking, but future studies are needed to determine whether our current findings are due to the activation of VP-associated circuitry or another basal forebrain area.

Dopaminergic innervation of the nucleus accumbens (**NAc**) has similarly been ascribed as having a causal role in sign-tracking, in which dopamine release was correlated with sign-tracking responses and dopamine receptor antagonists selectively reduced the acquisition of sign-tracking behavior (Danna and Elmer, 2010; Flagel et al., 2011; Fraser et al., 2016). Given the connectivity between the NAc and basal forebrain areas including the HDP and VP, it is likely that there is an interplay between dopaminergic and cholinergic neurotransmission within a circuit that promotes sign-tracking behavior. For example, one study showed that sign-trackers had increased dopaminergic activity while goal-trackers showed higher cholinergic activity in the cortex during presentations of a cue previously paired with cocaine (Pitchers et al., 2017a; 2017b). While this finding may be specific to a cocaine-predictive reward, it is one piece of evidence suggesting that these neurotransmitters systems contribute to different aspects of the sign-tracking response. More studies measuring the cholinergic response during the PavCA procedure are needed. Cholinergic signaling from the basal forebrain can modulate NAc dopamine release via nicotinic receptors on dopamine terminals (Threlfell et al., 2012). Thus, activation of basal forebrain cholinergic neurons might indirectly boost cue-evoked dopamine, enhancing the attribution of incentive salience. However, Threlfell manipulated cholinergic neurons within the NAcc, additional studies are needed to determine whether this is the case for cholinergic input arising from basal forebrain neurons in intact animals.

It is important to acknowledge that only male rats were used in this study. We and others have found that females tend to sign-track more than males (King et al., 2021; Hakus et al., 2025; Hughson et al., 2019, but see Bien and Smith, 2023). We chose males because we hypothesized an increase in sign-tracking, that we surmised would be difficult to detect in rats with already high levels of sign-tracking (i.e., females). Further, the PavCA paradigm was designed to promote behavioral variability and is vulnerable to cohort effects (King et al. 2021; Fitzpatrick et al., 2013), thus requiring large sample sizes to detect group differences. That said future studies should determine whether cholinergic stimulation would lead to even higher levels of sign-tracking or a ceiling effect.

## Conclusions

Overall, the current study provides evidence that the basal forebrain cholinergic system can promote the incentive salience attribution to reward cues. This has important implications for understanding how this system contributes to maladaptive cue-driven behaviors that underlie substance use disorders and other compulsive conditions in humans. In clinical populations, heightened reactivity to drug cues is a major factor in craving and relapse, and this cue-reactivity is often resistant to cognitive regulation (Childress et al., 1999; Volkow et al., 2008; 2010; Guerin et al., 2021). Thus, these findings raise the possibility that individual difference in basal forebrain cholinergic tone, can influence attentional biases and cue-reactivity in humans, thus contributing to vulnerability to compulsive reward seeking. This may lead to behavioral and pharmacological interventions that focus on managing cue salience before problematic reward-seeking is established.

## ACKNOWLEDGEMENTS AND DECLARATIONS

Research reported in this publication was supported by grants from the National Institute on Alcohol Abuse and Alcoholism (AA024112 and AA007583), and the National Institute on Drug Abuse (DA037844). None of the authors have any other disclosures.

## Notes

### Competing Interest Statement

The authors have declared no competing interest.

